# Mitochondrial dysfunction underlies monocyte immune deficiency in patients with severe alcohol-related hepatitis

**DOI:** 10.1101/2025.01.28.634915

**Authors:** Paul K. Middleton, Ricardo J. Aramayo, Alex Montoya, Pavel V. Shliaha, Simon Pope, Louise Fets, Nikhil Vergis, Mark R. Thursz, Helena M. Cochemé

## Abstract

Severe alcohol-related hepatitis (sAH) is a life-threatening form of alcohol-related liver disease (ARLD) associated with a significant short-term mortality. Opportunistic infections due to impaired immune function are a major cause of patient mortality. Mitochondrial dysfunction within the liver is a well-recognised feature of ARLD and sAH. However, whether these hepatic mitochondrial defects extend to the immune system of sAH patients, underlying their immune dysfunction, remains unclear. Here, we demonstrate that sAH monocytes exhibit an increased content of inefficient, dysfunctional mitochondria. These changes were underpinned by abnormal mitochondrial cristae ultrastructure, which were associated with depletion of cristae structural proteins and alterations in cardiolipin profiles. Overall, our study uncovers novel structural and functional mitochondrial defects, which likely contribute to impaired monocyte immune function in sAH.

## Introduction

Liver disease poses a significant global health challenge, accounting for 2 million deaths per year^(1)^. Within Europe, the rates of liver-related mortality have dramatically increased over the last several decades^(2, 3)^. Alcohol remains the leading cause of liver disease in Europe, and is attributed to 75% of liver-related deaths in the United Kingdom^(2, 3)^.

Alcohol-related liver disease (ARLD) refers to a spectrum of progressive alcohol-related liver damage that can take years to decades of alcohol misuse to culminate in irreversible scarring termed cirrhosis. However, at any point within this spectrum of disease, patients can develop an acute life-threatening syndrome termed alcohol-related hepatitis (AH)^(4)^.

AH is characterised by an acute immune-mediated inflammation of the liver associated with progressive jaundice and coagulopathy in patients with significant alcohol excess^(5)^. Importantly, AH is associated with a high short-term mortality, with one meta-analysis reporting a pooled 28-day mortality of 25%^(6)^. Severe alcohol-related hepatitis (sAH) is traditionally defined as a Maddrey Discriminant Function (MDF) ≥32^(7, 8)^. To reduce hepatic inflammation, prednisolone is currently the standard treatment for patients with sAH. However, corticosteroid use only shows a modest short-term mortality benefit, which does not persist at 90 days^(9, 10)^. Other interventions that have undergone randomised controlled trials (including pentoxifylline, infliximab, canakinumab, co-amoxiclav and anakinra) have not demonstrated benefit compared to placebo/prednisolone^(9, 11–15)^. Therefore, there remains a strong clinical need to identify disease-modifying therapies for sAH.

A major cause of mortality in patients with sAH, and a key limitation on prednisolone use, is the development of opportunistic infections. Patients with sAH have a >2-fold increased risk of infection compared to patients with decompensated ARLD cirrhosis^(16)^. The development of infection is associated with a significantly greater mortality risk^(17, 18)^. A major contributor to this susceptibility is an associated immune dysfunction. Monocytes are a key member of the innate immune system. Previous studies have identified impairments in the oxidative burst and cytokine production of sAH monocytes, which correlated to the risk of subsequent infection^(19, 20)^. However, the exact mechanisms underlying this immune dysfunction in sAH are not fully understood.

Over the last decade, the role of immune cell metabolism in regulating immune cell function has gained growing recognition, establishing the field of immunometabolism. Mitochondria in particular have emerged as key regulators of immune cell function^(21, 22)^. In murine macrophages, accumulation of succinate, mitochondrial membrane hyperpolarisation, and mitochondrial reactive oxygen species (mtROS) production are vital for lipopolysaccharide (LPS)-induced interleukin-1β (IL-1β) production^(23)^. Conversely, increased α-ketoglutarate (αKG) production and increased mitochondrial electron transport are implicated in the anti-inflammatory impact of interleukin-4 (IL-4) on murine macrophages^(24)^.

Hepatic mitochondrial dysfunction is a well-recognised feature of sAH. Explanted livers from patients with sAH are reported to have decreased mitochondrial content with altered morphology and increased mtDNA oxidative damage^(25)^. Furthermore, mitochondrial electron transport chain (ETC) activity was found to be decreased in liver biopsies from patients with sAH, with lower Complex I and III activity correlating with increased mortality^(26)^. However, whether these hepatic mitochondrial defects extend to the immune system has not been well explored. This study investigates and characterises mitochondrial dysfunction within circulating monocytes from patients with sAH.

## Results

### sAH monocytes have increased mitochondrial content with altered morphology

Utilising flow cytometry, we first assessed mitochondrial content in monocytes via MitoTracker Green staining of freshly isolated leukocytes from either patients with sAH or healthy volunteers (HV). sAH monocytes exhibited significantly increased MitoTracker Green fluorescence compared to HV (Fig. S1A, Fig.1A,B). To further investigate this finding, we applied transmission electron microscopy (TEM) as an orthogonal approach. Our TEM analysis corroborated the increased mitochondrial content in sAH monocytes (Fig. 1C,D), which was primarily driven by an increase in mitochondrial number (Fig. S1B,C).

**Figure 1.**
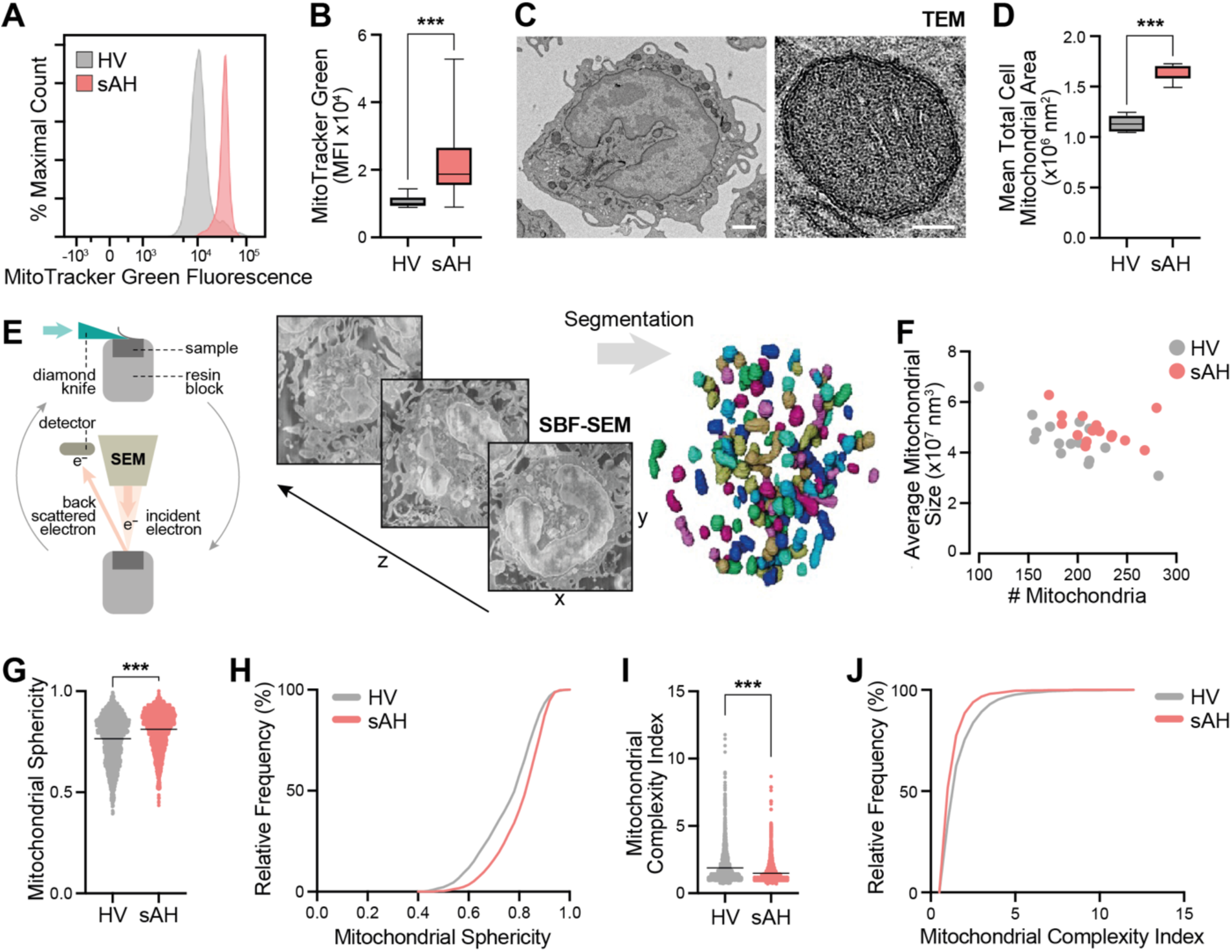
sAH monocytes have increased mitochondrial content with decreased structural complexity. **A)** Representative histogram of flow cytometry analysis for Mitotracker Green fluorescence between monocytes from healthy volunteer (HV) and severe alcohol-related hepatitis (sAH). **B)** Median fluorescence intensity (MFI) of MitoTracker Green signal by flow cytometry in circulating monocytes from HV (n=9) and sAH (n=14). *** p<0.001 (Mann-Whitney test). **C)** Representative transmission electron microscopy (TEM) image of a stained monocyte (left, scale bar=1 µm), with a magnified mitochondrion (right, scale bar=100 nm). **D)** Mean total mitochondrial area of 25 randomly selected monocyte cross sections, for n=5 individuals per group. *** p<0.001 (unpaired two-tailed t-test). **E)** Schematic representing the process of serial block-face scanning electron microscopy (SBF-SEM). **F)** Plot of pooled monocyte average mitochondrial number per cell to the average mitochondrial volume (n=15, 5 monocytes from 3 donors). **G)** Comparison of mitochondrial sphericity between pooled monocyte mitochondria from HV (n=2,844) and sAH (n=3,282). *** p<0.001 (Mann-Whitney test). **H)** Cumulative frequency distribution of mitochondrial sphericity between HV and sAH. **I)** Comparison of mitochondrial complexity index between pooled mitochondria from HV (n= 2,844) and sAH (n=3,282) from 5 monocytes from 3 donors. *** p<0.001 (Mann-Whitney test). **J)** Cumulative frequency distribution of mitochondrial complexity index between HV and sAH.

Since mitochondria exist as complex 3D networks, and TEM only provides a 2D view, we next employed serial block-face scanning electron microscopy (SBF-SEM) to image mitochondrial morphology within entire monocytes (Fig. 1E). Consistent with our flow cytometry and TEM data, total mitochondrial volume by SBF-SEM was greater in sAH monocytes compared to HV (Fig. S1D). Our 3D analysis also identified variation in inter-monocyte mitochondrial number and average mitochondrial volume, likely representing differences in mitochondrial network fission/fusion states (Fig. 1F). This distribution was altered in sAH monocytes, suggesting an increase in both mitochondrial number and average mitochondrial volume (Fig. 1F).

Analysis of pooled individual mitochondrial dimensions revealed an increase in sAH mitochondrial volume (Fig. S1E,F), as well as an increase in mitochondrial sphericity and a decrease in mitochondrial complexity (Fig. 1G-J), without significant changes in mitochondrial length (Fig. S1G,H). Decreased mitochondrial complexity and increased sphericity have recently been reported in muscle biopsies from patients with mtDNA defects, where increased genetic burden was associated with a greater shift towards simple mitochondrial morphology^(27)^. Despite increased mitochondrial content in sAH monocytes, mtDNA copy number was not significantly different to HV (Fig. S1I). Overall, sAH monocytes have increased mitochondrial content, with less complex 3D networks.

### sAH monocyte mitochondria are dysfunctional with inefficient respiration, impaired polarisation, altered TCA cycle metabolic profiles, and increased mtROS production

To investigate mitochondrial function, we conducted high resolution respirometry on freshly isolated monocytes from patients with sAH. To minimise the potential impact of isolation, monocytes were purified using negative magnetic bead selection, and respirometry was performed on intact monocytes in human plasma-like media (HPLM) using mitochondrial inhibitors and uncouplers. In keeping with the increased mitochondrial content, sAH monocytes had elevated basal (routine) and maximal oxygen consumption, with an increase in proton leak-associated respiration (Fig. 2A). ATP-linked respiration was not significantly increased compared to HV (Fig. 2A). These findings were corroborated when examined using a substrate-based approach on permeabilised monocytes (Fig. S2A-C).

**Figure 2.**
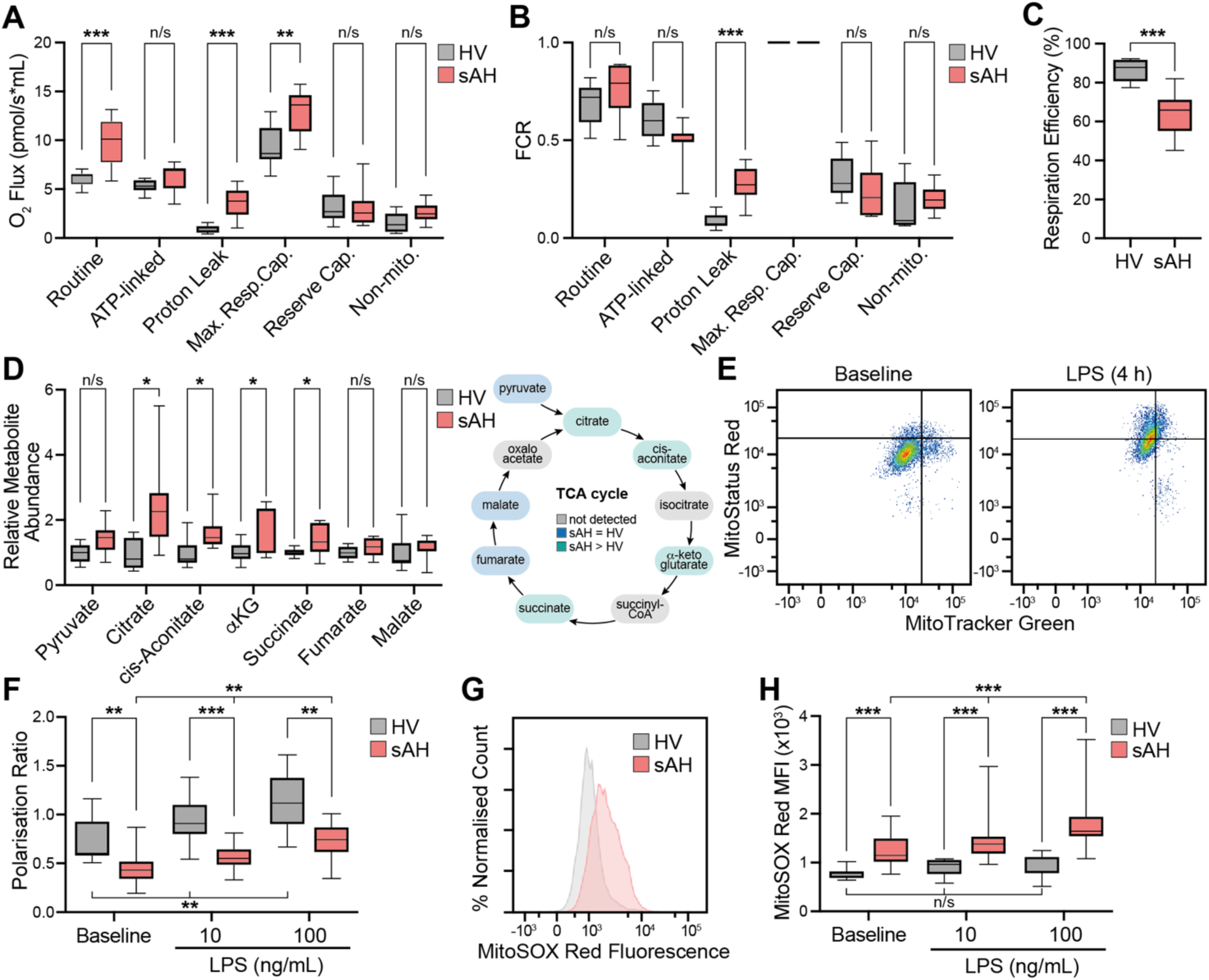
sAH monocytes have impaired mitochondrial respiratory efficiency, decreased membrane polarisation, altered TCA cycle profile, and elevated mtROS. A-B) High resolution respirometry of monocytes (1x10^6^ cells) from HV (n=8) and patients with sAH (n=10). O2 flux (**A**), and flux control ratio (FCR) of maximal respiratory capacity (**B**). ** p<0.01, *** p<0.001 (Mann-Whitney test). **C)** Mitochondrial respiratory efficiency. *** p<0.001 (unpaired two-tailed t-test). **D)** Left: Intracellular metabolomics of monocytes (1x10^6^ cells) from patients with sAH (n=7) and HV (n= 9). * p<0.05 (Mann-Whitney test). Right: Schematic of the tricarboxylic acid (TCA) cycle. αKG, α-ketoglutarate. Colour-coding: grey, not detected; blue, detected and not significantly different; green, detected and increased in sAH. **E)** Representative scatter plot of HV monocyte MitoTracker Green and MitoStatus Red staining at baseline and after lipopolysaccharide (LPS) stimulation (4 h, 100 ng/mL). **F)** Mitochondrial membrane polarisation at baseline and after the indicated LPS exposure (4 h). HV *vs* sAH pairwise comparisons: ** p<0.01, *** p<0.001 (Mann-Whitney test); LPS dose effect: ** p<0.01 (Friedman test). **G)** Representative histogram of flow cytometry analysis for MitoSox Red fluorescence. **H)** Monocyte mitochondrial superoxide production as measured by median fluorescence intensity (MFI) of MitoSOX Red at baseline and after 4 h of LPS stimulation (HV n=7, sAH n=14). HV *vs* sAH pairwise comparisons: *** p<0.001 (Mann-Whitney test); LPS dose effect: n/s p>0.05, *** p<0.001 (Friedman test).

To account for variations in mitochondrial content, respirometry values were normalised to maximal respiratory capacity (Fig. 2B). Interestingly, after normalisation, only proton leak-associated respiration remained significantly elevated in sAH monocytes (Fig. 2B). This increased proton leak resulted in significantly decreased mitochondrial coupling efficiency in sAH monocytes (Fig. 2C). Loss of mitochondrial coupling efficiency trended towards correlation with markers of liver dysfunction in sAH patients, including bilirubin levels and MELD score (Fig. S2D,E).

Intracellular metabolomics revealed altered tricarboxylic acid (TCA) cycle function with significant increased levels of citrate, cis-aconitate, α-KG and succinate in sAH monocytes compared to HV. However, fumarate and malate were not found to be significantly increased suggesting alterations in TCA cycle flux (Fig. 2D).

Given the increased proton-leak associated respiration, we assessed mitochondrial membrane polarisation using the membrane potential-dependent dye MitoStatus Red. MitoStatus Red fluorescence was normalised to the MitoTracker Green signal to account for observed variations in mitochondrial content. Stimulation with LPS lead to an increase in mitochondrial polarisation in both HV and sAH monocytes (Fig. 2E), as previously described for murine macrophages^(23)^. However, consistent with our finding of increased proton leak, sAH monocyte mitochondria had decreased polarisation at baseline and after LPS stimulation compared to HV (Fig. 2F).

We next assessed mtROS production using the superoxide-selective dye MitoSOX Red. sAH monocytes had significantly greater mtROS production at baseline compared to HV (Fig. 2G,H). Furthermore, in sAH monocytes, LPS stimulation was associated with a significant increase in mtROS production, which did not occur in HV monocytes (Fig. 2G,H). Altogether, sAH monocyte mitochondria are dysfunctional with inefficient respiration, impaired polarisation, altered TCA metabolic profiles, and increased mtROS production.

### sAH monocyte mitochondria display abnormal cristae ultrastructure, associated with depletion of MICOS proteins and altered cardiolipin profiles

To investigate the mechanisms underlying these features of mitochondrial dysfunction, we assessed detailed cristae ultrastructure by TEM. Interestingly, sAH mitochondria displayed structurally abnormal, swollen cristae (Fig. 3A). Maximal mitochondrial cristae width was significantly increased in sAH monocytes compared to HV (Fig. 3B,C). Untargeted proteomics of sAH monocytes revealed a significant depletion of proteins involved in maintaining cristae ultrastructure compared to HV, including subunits of the mitochondrial contact site and cristae organising system (MICOS) complex (MIC27, MIC60 and MIC19), as well as OPA1, another component of the mitochondrial inner membrane that regulates cristae junctions (Fig. 3D). Loss of MIC27 in sAH monocytes was independently validated by western blotting on a separate cohort of patient samples (Fig. 3E,F). Levels of other mitochondrial proteins, such as the MTCO2 subunit of Complex IV, were unaffected (Fig. 3E,F).

**Figure 3.**
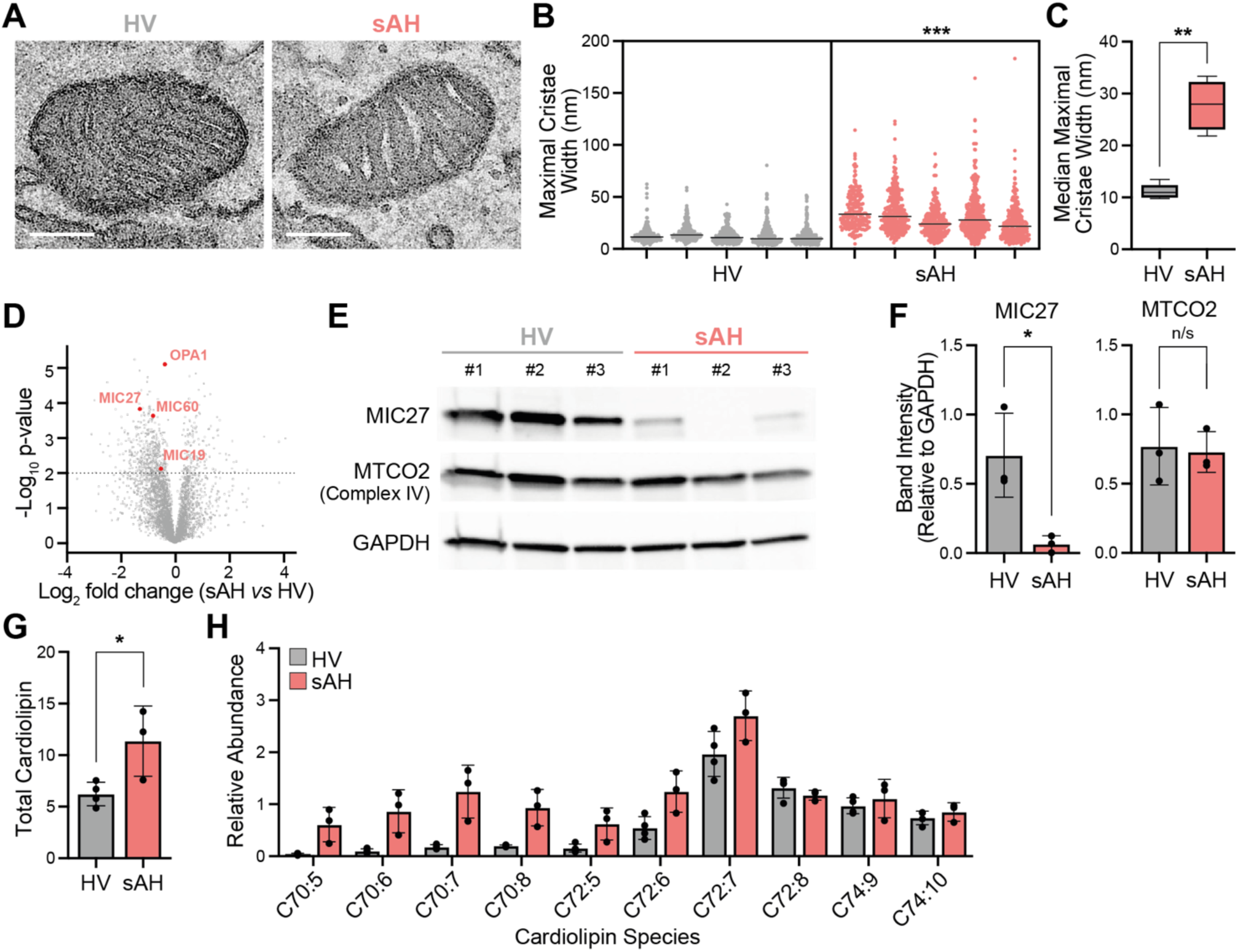
sAH monocytes have abnormal mitochondrial cristae ultrastructure and altered cardiolipin profiles. **A)** Representative TEM images of a mitochondrion within a monocyte from HV (left) and a patient with sAH (right). Scale bar=200 nm. **B)** Nested plot of individual maximal cristae width per mitochondria from 25 monocytes (n=5 individuals per condition). *** p<0.05 (nested t-test). **C)** Median maximal cristae width per individual (n=5). ** p<0.01 (Mann-Whitney test). **D)** Untargeted proteomics from monocyte protein extracts of HV (n=5) and sAH (n=4). Dotted line denotes level of significance (adjusted p value <0.05). **E)** Western blot of monocyte protein samples against MIC27, MTCO2 (subunit of Complex IV) and GAPDH (n=3 individuals per condition). **F)** Quantification of band intensity relative to GAPDH. * p<0.05 (unpaired two-tailed t-test). **G)** Quantification of total cardiolipin by mass spectrometry within lipids extracted from 1x10^6^ monocytes, comparing HV (n=4) and sAH (n=3). * p<0.05 (unpaired two-tailed t-test). **H)** Cardiolipin species, plotted by total acyl chain carbon content and number of double bonds (e.g. C72:8 = cardiolipin with 4x C18:2 fatty acyl chains).

Mitochondrial membrane phospholipids, such as cardiolipin, are another key determinant of cristae structure^(28)^. Cardiolipin was analysed on lipids extracted from purified monocytes by LC-MS. Consistent with our previous findings of increased mitochondrial content, total cardiolipin levels were increased in sAH monocytes (Fig. 3G). Interestingly, detailed cardiolipin profiling of sAH monocytes indicated an increase in shorter-chained (C70) species (Fig. 3H). Altogether, these data show that monocyte mitochondria have disrupted cristae ultrastructure with depletion of cristae structural proteins and altered cardiolipin profiles.

### Circulating sAH monocytes have impaired LPS-stimulated cytokine production

Given our findings of mitochondrial dysfunction, we next sought to characterise the immune phenotype and function of sAH monocytes. Analysis of circulating monocytes in sAH patients identified an increase in intermediate monocytes (CD14hi CD16+), with a relative decrease in classical (CD14hi CD16-) and non-classical (CD14lo CD16+) sub-types (Fig. 4A). Increased intermediate monocytes have frequently been reported in conditions associated with immune activation, including rheumatoid arthritis, atherosclerosis and sepsis^(29–31)^. Decreased HLA-DR expression is associated with impaired monocyte function and is predictive of mortality in severe sepsis^(32, 33)^. While at baseline there was a trend towards decreased HLA-DR expression in unstimulated sAH monocytes, which did not reach statistical significance, LPS-induced HLA-DR expression was significantly decreased in sAH compared to HV monocytes (Fig. 4B). Decreased LPS-induced HLA-DR expression has previously been reported to differentiate survivors from non-survivors in patients with severe sepsis^(34)^.

**Figure 4.**
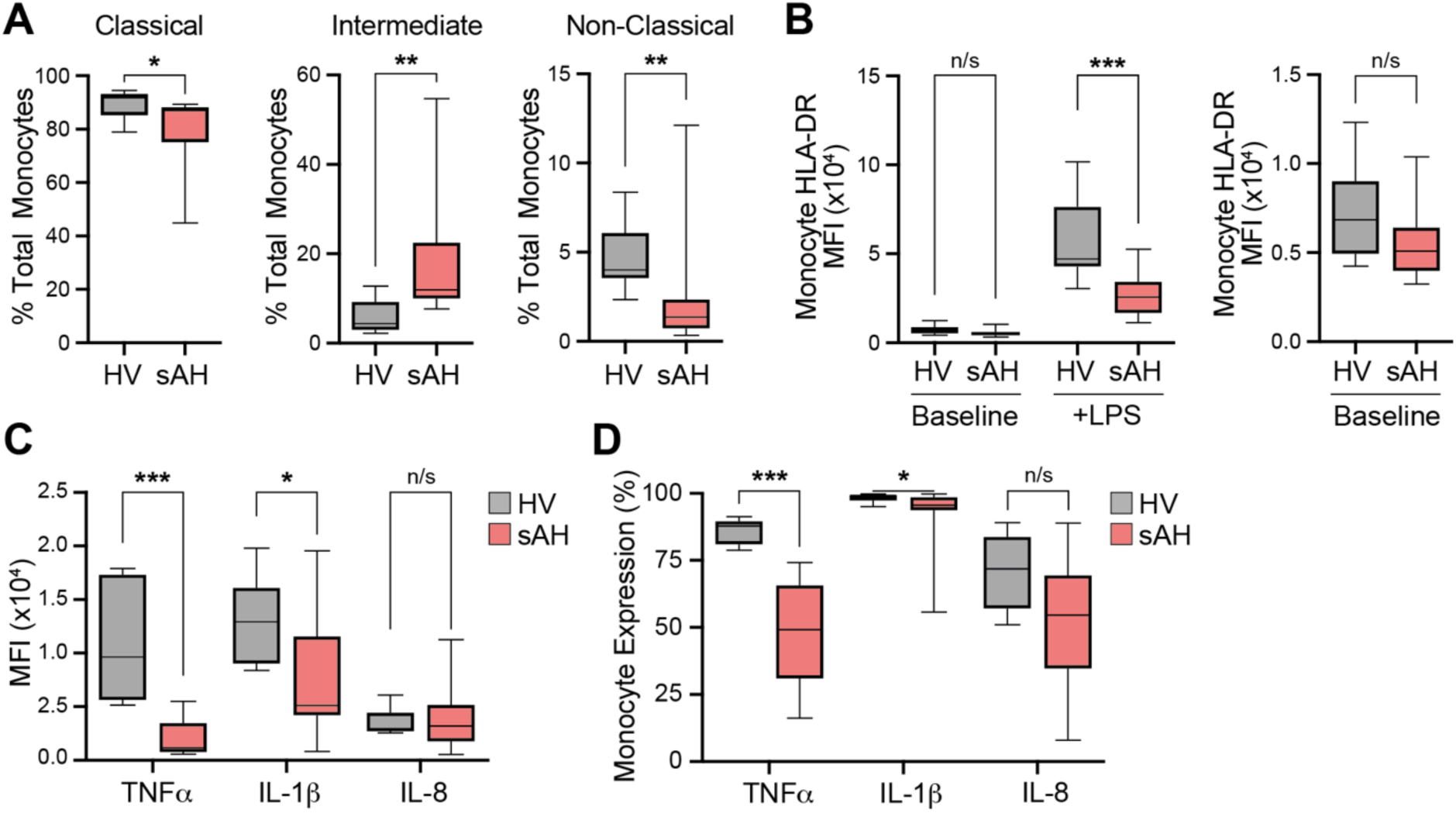
Circulating monocytes have impaired LPS-stimulated cytokine production. **A)** Circulating monocyte sub-sets expressed as a proportion of total monocytes from HV (n=8) and sAH patients (n=14). * p<0.05, **p<0.01 (Mann-Whitney test). **B)** Baseline and LPS-stimulated monocyte HLA-DR expression plotted as MFI (median fluorescence intensity). n/s p>0.05, *** p<0.001 (Mann-Whitney test). **C)** LPS-induced monocyte cytokine production expressed as MFI. **D)** Cytokine staining expressed as percentage positivity. n/s p>0.05,* p<0.05, *** p<0.001 (Mann-Whitney test). TNFα, tumour necrosis factor α; IL-1β, interleukin-1β; IL-8, interleukin 8.

Finally, LPS-induced cytokine production was assessed using intracellular cytokine staining. Strikingly, LPS-induced TNFα and IL-1β production were significantly impaired in sAH monocytes compared to HV, with no significant change in IL-8 (Fig. 4C,D). Overall, these results show that sAH monocytes have impaired immune function.

## Discussion

This study has revealed previously uncharacterised structural and functional mitochondrial defects in sAH monocytes. sAH monocytes displayed an increased quantity of inefficient, dysfunctional mitochondria, resulting in increased mitochondrial oxygen consumption and cellular mtROS. Our work has also confirmed impaired pro-inflammatory cytokine production in sAH monocytes^(20)^. These mitochondrial defects may underlie the monocyte immune dysfunction in sAH patients.

### Mitochondria and Monocyte Immune Function

Increased monocyte mitochondrial content has been reported in patients with sepsis requiring intensive care^(35)^. Similar to sAH, monocytes from patients with severe sepsis exhibited impaired LPS-induced cytokine production^(36)^. Despite an increase in mitochondrial content of sAH monocytes, we did not identify a corresponding increase in mtDNA copy number. Intriguingly, this phenotype has previously been described in murine macrophages induced into LPS tolerance by either mitochondrial stress or LPS pre-exposure^(37)^. Therefore, mtDNA/mitochondrial mass disequilibrium may be an interesting future avenue of investigation.

Mitochondrial content is regulated by mitochondrial biogenesis and mitophagy. Peroxisome proliferator-activated receptor gamma (PPARγ) is a transcription factor which promotes mitochondrial biogenesis^(38)^. PPARG expression was been found to correlate to anti-inflammatory marker expression on human macrophages^(39)^. Furthermore, PPARG agonists promoted differentiation of monocytes into anti-inflammatory macrophages^(39)^. Interestingly, PPARG transcription was found to be significantly upregulated in sAH monocytes^(20)^.

Our finding of increased mitochondrial oxygen consumption in sAH monocytes is contrary to a recent study which reported decreased mitochondrial oxygen consumption in sAH PBMCs (peripheral blood mononuclear cells)^(40)^. However, this discrepancy may be explained by the heterogeneous nature of PBMCs. Indeed, previous studies have shown poor correlation between mitochondrial functional analysis of PBMCs and PBMC sub-groups, including monocytes^(41)^. A major contributor to this is the contamination of PBMCs with platelets, which contain mitochondria^(41)^. Given thrombocytopenia is a common finding in advanced chronic liver disease, this is may be a significant confounding factor. Therefore, the analysis of purified monocytes, rather than bulk PBMCs, is a significant advantage of our study.

Increased mitochondrial oxygen consumption is also reported in metabolic dysfunction-associated steatohepatitis (MASH) monocytes, along with increased TNFα and IL-1β production^(42)^. This elevated cytokine phenotype was recapitulated in HV monocytes by LPS stimulation^(42)^. However, unlike in sAH monocytes, the increased mitochondrial respiration in MASH and LPS-stimulated monocytes was accompanied by a proportionate and significant increase in ATP-linked respiration, suggesting that mitochondrial efficiency was maintained^(42)^. Therefore, our observation of mitochondrial inefficiency may be an important factor in sAH monocyte dysfunction. LPS is known to be elevated in the serum of patients with sAH^(43)^, which could promote changes in monocyte mitochondrial function.

Consistent with increased mitochondrial mass and activity, we identified an increase in TCA cycle metabolites, including citrate, cis-aconitate, α-KG and succinate. Interestingly, other TCA metabolites from the latter half of the cycle (e.g. fumarate and malate) were not significantly raised. This pattern differs from a previous study examining intracellular metabolite changes in LPS-stimulated healthy monocytes, which identified significant increases in malate and fumarate, but reported no significant change in citrate, succinate or α-KG^(44)^. Altered TCA cycle metabolism may underlie changes in monocyte immune function. Pre-treatment with exogenous fumarate has been shown to promote *C. albicans*-induced TNFα production in healthy monocytes, suggesting increased fumarate may contribute to monocyte immune response^(45)^. Similarly, in murine macrophages fumarate accumulation was found to promote pro-inflammatory cytokine production^(46)^. Conversely, increased α-KG was shown to be critical for the anti-inflammatory response of macrophages to interleukin-4 (IL-4)^(24)^. Altered TCA cycle metabolites may suggest compensatory changes in metabolism in response to impaired mitochondrial efficiency. Glutaminolysis, for example, has been shown to increase in response to mtDNA defects^(47)^.

Although we identified an increase in succinate accumulation, IL-1β production was significantly impaired contrary to previous findings in murine macrophages^(23)^. This may be due to differences in mitochondrial membrane polarisation. As in murine macrophages, HV monocytes exhibited mitochondrial membrane hyperpolarisation in response to LPS. Conversely, sAH monocytes were relatively depolarised at baseline, and had significantly decreased polarisation after LPS stimulation.

In keeping with mitochondrial dysfunction, mtROS was significantly increased in sAH monocytes. Furthermore, stimulation with LPS was associated with a significant increase in mtROS. mtROS have previously been implicated in pro-inflammatory immune responses in macrophages and monocytes^(23, 48, 49)^. However, prolonged increase in mtROS in murine macrophages is associated with an increase in mitochondrial chaperones and antioxidant enzymes, which are thought to contribute to the development of LPS tolerance^(37)^.

### Cristae Structure and Mitochondrial Function

We identified significant disruption of the cristae ultrastructure in sAH monocytes, which was associated with depletion of cristae structural proteins including OPA1, and the MICOS components MIC27, MIC60 and MIC19, as well as changes in the profile of cardiolipin species. Cristae structure influences mitochondrial function through several mechanisms. Disruption of cristae shape, through depletion of OPA1, can impair ETC supercomplex formation^(50)^. ETC supercomplexes are recognised to maintain efficient transfer of electrons between complexes and decrease mtROS production^(50–52)^. Defective supercomplex formation may underlie the respiratory inefficiency and increased mtROS production. The mitochondrial cristae have also been found to retain their own electrochemical gradients separate from the intermembrane space, allowing each cristae to function independently similar to cells within a battery^(53)^. Disruption of cristae structure through depletion of MIC60 or OPA1 can dissipate the cristae electrochemical gradient^(53)^. Therefore, defects in cristae ultrastructure may lead to decreased mitochondrial polarisation and respiratory efficiency.

Additionally, the cristae junction is proposed to regulate mitochondrial matrix calcium concentration by controlling access to calcium transporters in the cristae^(54)^. The cristae junction is maintained by the MICOS complex as well as OPA1^(54)^. Knockdown of OPA1 in mitochondria leads to increased calcium uptake^(55)^. Calcium promotes the activity of key TCA cycle enzymes including pyruvate dehydrogenase, isocitrate dehydrogenase and α-ketoglutarate dehydrogenase^(56)^. Therefore, altered cristae structure may underlie increased mitochondrial respiration and altered TCA cycle activity.

Altogether, our study uncovers novel morphological and functional features of mitochondrial dysfunction in sAH monocytes, which are likely underpinned by disruption of cristae ultrastructure. Future work will focus on characterising immunometabolic pathways linking these mitochondrial changes to impaired monocyte immune function, with the aim of uncovering novel therapeutic targets.

## Methods

### Participant Characteristics

Patients with a clinical diagnosis of severe alcohol-related hepatitis (sAH) were recruited to clinical studies with ethical approval for immunological research including Multicentre Cohort Study in Alcoholic Hepatitis (MICAH) [IRAS ID 258447], and Prospective Investigation of Steatohepatitis and Scaring in Heavy Alcohol Drinkers (ALCOBASE) [IRAS ID 202572]. All patients conformed to the inclusion criteria for patients with alcohol-related hepatitis as outlined by The National Institute on Alcohol Abuse and Alcoholism Alcoholic Hepatitis Consortia^(8)^. Alternative causes of liver dysfunction including viral hepatitis, biliary obstruction and hepatocellular carcinoma were excluded. Patients with concomitant liver disease including hepatitis B, hepatitis C, autoimmune liver disease, Wilsons disease, drug-induced liver injury, haemochromatosis or hepatocellular carcinoma were not included. sAH was defined as a Maddrey Discriminant Function (MDF) of ≥32. All samples were collected prior to commencement of prednisolone or investigative drug. Healthy volunteers (HV), who drank less than 14 units of alcohol per week, were recruited to clinical studies with ethical approval for immunological research including ALCOBASE [IRAS ID 202572] and Monocytes and Macrophage Study [IRAS 87203]. Baseline characteristics of participants who contributed samples to this research are described in Table 1.

**Table 1.** Summary of participant demographics. Characteristics of participants with severe alcohol-related hepatitis (sAH) and healthy volunteers (HV). Mean values with range in parentheses (^#^, unpaired t-test; ^##^, chi-square test).

|  | sAH (n=35) | HV (n=20) | p value |
| --- | --- | --- | --- |
| Age (years) | 48 (30-66) | 34 (24-60) | <0.001<sup>#</sup> |
| Sex, male (%) | 66 [24/35] | 75 [15/20] | n/s<sup>##</sup> |
| Total Bilirubin (μmol/L) | 280 (70-677) | - | - |
| Albumin (g/L) | 25 (16-45) | - | - |
| INR | 1.5 (1.2-5.5) | - | - |
| ALT (U/L) | 36 (6-125) | - | - |
| AST (U/L) | 118 (52-386) | - | - |
| Creatinine (μmol/L) | 61 (43-148) | - | - |
| Maddrey Discriminant Function (MDF) | 54.4 (37.8-234.2) | - | - |
| MELD score | 22 (16-38) | - | - |

### Flow Cytometry

100 μL of unstimulated or LPS-stimulated whole blood was separated into a FACS tube per experiment. Red blood cells (RBC) were lysed using RBC lysis buffer (eBioscience). The leukocyte pellet was then resuspended in an antibody cocktail dependant on experimental plan (Supplementary Table 1) and incubated for 20 min at room temperature. MitoTracker Green FM (20 nM; Invitrogen), MitoStatus Red (1 nM; BD Biosciences) and MitoSOX Red (1 μM; Invitrogen) were used to quantify mitochondrial content, polarisation and mtROS respectively. LPS stimulation was performed on 1 mL of whole blood at 37°C for 4 h in the presence of 10 ng/mL or 100 ng/mL LPS derived from *Escherichia coli* (*E. coli*) O111:B4 (Sigma Aldrich).

Intracellular cytokine staining was performed by stimulation of 1 mL of whole blood with 100 ng/mL LPS in the presence of Protein Transport Inhibitor Cocktail (eBioscience) for 4 h at 37°C. Fixation and permeablisation were performed using the Intracellular Fixation & Permeabilization Buffer kit (eBioscience) as per manufacturer instructions. Cells were incubated with an antibody cocktail to surface markers and intracellular cytokines at 4°C (Supplementary Table 2).

Flow cytometry was performed on a 3 laser LSRFortessa Cell Analyzer (BD Biosciences), and analysed using FlowJo software (version 10.8). Monocytes were identified as Fixed Viability Dye 450-/CD3-/CD19-/CD56-/CD11b+/HLADR+/CD15- cells. These cells were then gated by CD14 and CD16 positivity to sub-classify classical (CD14hi CD16-), intermediate (CD14hi CD16+) and non-classical (CD14lo CD16+). Total monocytes gating included all sub-groups (Fig. S1A). Fluorescence was quantified as median fluorescence intensity (MFI). Polarisation ratio was calculated by dividing MitoStatus Red MFI by MitoTracker Green MFI to normalise for variations in mitochondrial content. For intracellular cytokine staining, positivity gating was determined using a fluorescence minus one gating strategy, where gating was determined using a control sample lacking the fluorophore of interest.

### Monocyte Isolation

PBMCs (peripheral blood mononuclear cells) were separated from whole blood by density centrifugation using Ficoll-Paque Premium 1.073 (Cytiva). Monocytes were isolated from PBMCs with negative magnetic bead selection using the Pan Monocyte Isolation Kit (Miltenyi Biotec) as per manufacturer instructions.

### Electron Microscopy

Freshly isolated monocytes were fixed and stained using a modified NCMIR Method for 3D electron microscopy^(57)^. Samples were embedded in epoxy embedding medium (Sigma Aldrich).

Transmission electron microscopy (TEM) was performed on 50 nm sections of sample, processed using a Leica EM UC7 Ultramicrotome, and floated on top of 300 mesh copper grids with a thin continuous carbon layer. Images were recorded on a Thermo Fisher Talos F200i electron microscope, operated at 200 kV acceleration voltage, using a Falcon 3EC direct electron detector operating in integration mode. 25 randomly selected monocytes were imaged per sample. The monocyte cross-sections were imaged at x5,300 magnification, with a final pixel size of 2.72 nm using a -15 μm defocus. The total electron dose was 32 e^−^/nm^2^. Individual mitochondria were then imaged at x28,000 magnification, with a final pixel size of 0.52 nm using a -10 μm defocus. The total electron dose was 838 e^−^/nm^2^. Image analysis was performed using the open source software Fiji (version 1.54)^(58)^. Each mitochondrion was outlined and the dimensions recorded including area, perimeter and Feret diameter. Maximal cristae width was quantified by measuring the maximal intracristae distance (perpendicular to the cristae direction) within each mitochondrion.

For serial block-face scanning electron microscopy (SBF-SEM), resin-embedded samples were sputter coated with a 14 nm-thin layer of gold using an EM ACE600 High Vacuum Sputter Coater (Leica). SBF-SEM was performed using the Apreo VolumeScope (Thermo Fisher). The sample was imaged in low vacuum mode with a beam energy of 4.5 keV and the VS-DBS detector. Image acquisition was performed using Maps software (version 3.9). A sample area of 20.48 μm^2^ was selected. Imaging was performed using a 300 ns dwell time and 4 frame integration to a final pixel size of 3.33 nm. Sample cutting was performed at a depth of 50 nm. Automated image aligning was performed every 25 slices. Image analysis was performed using Amira software (version 2020.3, Thermo Fisher). From SBF-SEM volumes, 5 entire monocytes per individual were selected for detailed analysis. Images were de-noised using the Amira Backscatter Electron Scanning Electron Microscope U-net denoiser module. Individual mitochondria were then manually segmented. Mitochondrion dimensions including surface area (Area3D), volume (Volume3D) and length (Length3D) were quantified. Mitochondrial complexity index and sphericity were calculated following published equations^(27, 59)^.

### Mitochondrial DNA copy number

DNA was extracted from isolated monocytes using the Puregene DNA isolation kit according to manufacturer instructions (Qiagen). DNA concentration was determined using a NanoDrop One spectrophotometer (Thermo Scientific). DNA samples were diluted to 1 ng/μL. Primers targeting human genomic DNA (β2 microglobulin; F: TGCTGTCTCCATGTTTGATGTATCT, R: TCTCTGCTCCCCACCTCTAAGT) and mtDNA (mitochondrial tRNALeu(UUR); F: CACCCAAGAACAGGGTTTGT, R: TGGCCATGGGTATGTTGTTA) were used to quantify nuclear and mitochondrial genome content via quantitative polymerase chain reaction (qPCR) following a published equation^(60)^. Real time PCR was performed on a QuantStudio 7 Flex Real-Time PCR system (Thermo Fisher), and analysed using QuantStudio Design & Analysis Software (Thermo Fisher, version 2.5.0).

### High Resolution Respirometry

High resolution respirometry was carried on using the Oxygraph O2k Respirometer, fitted with the 0.5 mL small volume chamber (Oroboros Instruments). Oxygen flux was corrected to background flux over a range of oxygen concentrations following a dithionite titration as described by the manufacturer. 1 x 10^6^ monocytes were used for each experiment in both Chamber A and Chamber B, allowing for two technical replicates per sample. A median of technical replicates was used in data analysis.

Respirometry was performed on intact monocytes using HPLM (Gibco). Briefly, oxygen consumption was allowed to stabilise for 20 min, then baseline respiration (Routine) was measured. Next, 2 μL of 250 nM oligomycin (1 nM final; Sigma) was added to inhibit Complex V activity, and the sample was incubated for 20 min until oxygen consumption reached a steady state (Proton Leak). Serial 1 μL injections of 50 μM CCCP (500 nM steps; Abcam) were performed until maximal respiratory capacity (Max. Resp. Cap) was reached. Finally, 1 μL of 250 μM rotenone (500 nM final; Sigma) and 1 μL of 1.25 mM antimycin A (2.5 μM final; Sigma) were added to inhibit mitochondrial oxygen consumption (ROX, residual oxygen consumption;). During data analysis, ROX was subtracted from other respiratory states. ATP-linked respiration was calculated by subtracting Proton Leak respiration from Routine respiration. Spare respiratory capacity (Reserve Cap.) was calculated by subtracting routine respiration from maximal respiratory capacity. Flux control ratio (FCR) of maximal respiratory capacity was calculated by dividing all respiratory states by maximal respiratory capacity. Mitochondrial coupling efficiency was calculated by expressing ATP-linked oxygen consumption as a percentage of routine respiration.

Respirometry was performed on permeabilised monocytes using mitochondrial respiration medium (Mir05; Oroboros Instruments). Briefly, 1 x 10^6^ monocytes were added to each chamber and the chambers closed. Oxygen consumption was allowed to stabilise for 20 min then then baseline respiration (ROUTINE) was measured. 1 μL of 2.5 mg/mL digitonin (Sigma) in dimethyl sulfoxide (DMSO; Sigma) was added to permeabilise the cell membrane. This dosage of digitonin was optimised through prior increasing titration of digitonin in the presence of succinate and adenosine diphosphate (ADP; Sigma). 1 μL of 5 mM cytochrome c (10 μM final; Sigma) followed by 1 μL of 2.5 M pyruvate (5 mM final; Sigma) and 2.5 μL of 50 mM malate (100 μM final; Sigma). Oxygen consumption in this state is due to proton leak, slip and cation cycling (State 2). 5 μL of 500 mM ADP (1 mM final; Sigma) with 300 mM magnesium chloride (600 μM final; Sigma) was then added. Oxygen consumption in this state is attributed to oxidative phosphorylation (State 3) Oxidative phosphorylation capacity (OXPHOS) was calculated by subtraction of State 2 from State 3 respiration. 1 μL of 62.5 μM CCCP (125 nM final; Abcam) was added to determine if OXPHOS was limited by ATP synthase capacity. 2.5 μL of 2 M glutamate (10 mM final; Sigma) was then added to complete all NADH-linked substrates (Complex I-linked State 3u). 5 μl of 1 M succinate (10 mM final; Sigma) was added to provide a Complex II substrate (Complex I+II-linked State 3u). 1 μL of 250 μM rotenone (500 nM final; Sigma), a Complex I inhibitor, was added to measure oxygen flux of Complex II-linked respiration alone (Complex II-linked State 3u). 5 μL of 1 M glycerol-3-phosphate (10 mM final; Sigma) was added to provide electrons directly to ubiquinone in order to fuel the electron transport chain (ETC) at Complex III. 1 μL of 1.25 mM antimycin A (2.5 μM final; Sigma), a Complex III inhibitor, was added blocking ETC oxygen consumption leaving residual oxygen consumption (ROX). ROX was subtracted from flux in other respiratory states during data analysis.

### Intracellular Metabolomics

After isolation, monocytes were allowed to rest for 20 min at room temperature in HPLM. 1 x 10^6^ cells were then separated, washed with DPBS, centrifuged and the supernatant removed. The resulting cell pellet was flash frozen on dry ice and stored at -80°C until processing. For analysis, cell pellets were resuspended in a mixture of methanol, water and chloroform (725, 90 and 80 μL respectively) containing 1 nM *scyllo*-inositol. Samples underwent 3 rounds of sonication for 8 min at 4°C, and left to extract at 4°C until the end of the day. Samples were then centrifuged at 4°C on full speed for 5 min to pellet precipitated protein. The supernatant was removed and dried within a vacuum concentrator. Protein pellets were saved for future proteomic analysis. Biphasic extraction was carried out to separate polar and apolar metabolites using a chloroform, methanol and water solution (1:3:3 ratio). The dried supernatant was resuspended in 350 μL of this solution and vortexed to ensure mixing. This was then centrifuged to separate the phases. The aqueous phase containing polar metabolites was removed and dried within a glass vial insert. The non-aqueous phase containing non-polar metabolites and lipids was stored at -80°C for future lipid analysis. The dried polar metabolites were washed twice with 40 µL of methanol and dried again to minimise water content. The sample was then derivatised using 20 µL of 20 mg/mL methoxyamine in pyridine at room temperature overnight before addition of 20 µL of N,O-bis(trimethylsilyl)trifluoroacetamide with 1% trimethylchlorosilane (Sigma) for ≥1 h.

Metabolite analysis was performed via gas chromatography-mass spectroscopy (GC-MS) using the Agilent 8890B-5977A system. Splitless injection (injection temperature 270°C) onto a 30 m +10 m × 0.25 mm DB-5MS + DG column (Agilent J&W) was used, with a helium carrier gas, in electron-impact ionisation (EI) mode. Oven temperature was initially 70°C (2 min), and was subsequently raised to 295°C at 12.5°C/min, then to 320°C at 25°C/min (held for 3 min). MassHunter Workstation software (version B.06.00 SP01, Agilent Technologies) was used for metabolite identification and quantification by comparison to the retention times, mass spectra and responses of known amounts of authentic standards.

### Proteomics

Protein pellets were resolubilised using the previously described SEPOD surfactant cocktail containing 2% SDS, 1%SDC and 2% IGEPAL CA-630 and extracted at 70°C^(61)^. Protein concentration was quantified using the BCA assay. 50 μg of protein was then resuspended in 25 μL and mixed with an equal volume of an aqueous solution containing 20 mM TCEP and 40 mM chloroacetamide. The samples then underwent precipitated assisted capture^(62)^ with the addition of 100 µg of hydroxyl beads (MR-HYX010) and 75 µL ethanol. Beads were washed in 80% ethanol three times before undergoing overnight digestion in a 50 mM ammonium bicarbonate buffer containing trypsin (20 ng/µL) and LysC (10 ng/µL). The digest was transferred to a new plate and the resin was washed with 10 µL of 1% TFA and the washing solution was combined with the digest. LC-MS analysis was performed on an Ultimate 3000 RSLC nano liquid chromatography system (Thermo Scientific) coupled to a Q Exactive HF-X mass spectrometer (Thermo Scientific) via an EASY-Spray source. Electro-spray nebulisation was achieved by interfacing to Bruker PepSep emitters (PN: PSFSELJ20, 20 µm). Peptide solutions were injected directly onto the analytical column (self-packed column, CSH C18 1.7 µm beads, 300 μm × 35 cm) at a working flow rate of 5 μL/min for 4 min. Peptides were then separated using a stepped gradient of 0-45% buffer B over 66 min (composition of buffer A, 95/5%: H_2_O/DMSO + 0.1% FA; buffer B, 75/20/5% MeCN/H_2_O/DMSO + 0.1% FA), followed by column conditioning and equilibration. Eluted peptides were analysed by the mass spectrometer in positive polarity using a data-independent acquisition mode as follows: an initial MS1 scan was carried out at 120,000 resolution with an AGC target of 3e6 for a maximum IT of 200 ms, m/z range: 350-1650. This was followed by 30 DIA scans with variable window width at 30,000 resolution. AGC target set to 3e6 with maximum IT on auto. Normalised collision energy was set to 27%. Total run acquisition time was 82 min.

Data were processed using the Spectronaut software platform (Biognosys, version 17.2.230208.55965)^(63)^. Analysis was carried out in direct DIA mode as follows:

1. Pulsar Search: library generation and database search carried out using default settings for a tryspin/p specific digest as follows - missed cleavage rate set to 3 and variable modifications allowed for methionine oxidation, protein N-terminal acetylation, asparagine deamidation and cyclisation of glutamine to pyro-glutamate. PSM, Peptide and Protein group FDR = 0.01. Searches were carried out against the Uniprot Homo sapiens protein sequence database (downloaded 31/12/2023, 20,594 entries). Additionally, searches were also carried against a universal protein contaminants database (downloaded 04/06/2022, 381 entries)^(64)^.
2. Direct DIA+ (Deep) analysis: a mutated decoy database approach was employed with protein q-value cut-off for the experiment set to 0.01 at the identification level. Quantification set to MS2 with proteotypicity filtering by only proteotypic peptides with no value imputation strategy employed. Protein quantification method set to MaxLFQ^(65)^.

Both raw and normalised (automatic setting – local normalised) protein intensities were exported from Spectronaut for further processing. Exploratory and technical analysis of data carried out using in-house developed Python pipeline with various visualisations using both the Plotly and Pandas plotting libraries. Spectronaut protein level output tables were then filtered to remove hits to the contaminants database, before further analysis on normalised protein intensities.

Statistical testing was carried out using the Perseus platform (version 1.6.15.0)^(66)^. Human ontology annotations GOBP, GOMF, GOCC, KEGG and PROSITE (downloaded through the Perseus GO annotation tool) were assigned by Uniprot protein accessions. Data was log2 transformed before additional filtering and statistical testing. For multi-group comparisons, data was filtered for proteins with ≥4 replicate intensities per experimental group. Protein intensities of one-way ANOVA significant hits (multiple testing correction = permutation-based FDR, threshold = 0.05) were then z-scored in order to HCA-heatmap cluster the data. HCA row cluster IDs were assigned manually and subsequently used to carry out GO enrichment analysis (Fishers Exact test, multiple-testing correction = Benjamini-Hochberg, FDR threshold = 0.05). For 2-group comparisons, intensities were filtered as previously described and Student’s t-test carried out (multiple testing correction = permutation-based FDR, threshold = 0.05). t-test results were visualised as volcano plots. Intensities of significant hits were z-scored and further processed for GO enrichment analysis as previously described.

### Western Blotting

Isolated monocytes were spun to a pellet and solubilised with RIPA buffer (Thermo Fisher) supplemented with Halt Protease and Phosphatase inhibitor cocktail (Thermo Fisher). Cells were sonicated within a chilled water bath using a Diagenode biorupter (10 cycles of 30 s on/30 s off, low intensity) in order to sheer DNA and ensure complete protein solubilisation. Protein content was quantified using the Pierce BCA assay kit (Thermo Fisher), and 20 μg of protein was loaded per sample. Electrophoresis was performed on Criterion TGX Stain Free gels (Bio-Rad) using SDS buffer, and Transfer was performed with Bio-Rad Trans-blot Turbo onto nitrocellulose membrane (Bio-Rad). Primary antibodies included anti-MIC27 (anti-APOOL, Atlas Antibodies; HPA000612), anti-MTCO2 (Abcam; EPR3314) and anti-GAPDH (Sigma Aldrich; G9545). Anti-Rabbit HRP conjugate was used as a secondary antibody (Cell Signaling; 7074S). Chemiluminescence was elicited using SuperSignal West Pico Plus Chemiluminescent Substrate (Thermo Fisher). Membrane chemiluminescence was then visualised using a ChemiDoc Touch Imaging System (Bio-Rad), and quantified with Image Lab software (Bio-Rad, version 6.1.0).

### Cardiolipin Analysis

Lipid extracts were dried down under nitrogen and resuspended in 100 μL of methanol containing 0.4 μM of (C14:0)_4_ CL internal standard. 15 μL was injected into a Waters Acquity Ultra Performance Liquid Chromatography (UPLC) system (Waters Corporation). A gradient elution method was used as described^(67)^. An Acquity HSS T3 column was used (2.1 × 100 mm, 1.8 μm particle size; Waters). The column was maintained at 45°C. Mobile phase A was water/methanol = 6/4 + 10 mM ammonium formate + 0.1% FA and solvent B was isopropanol/methanol = 9/1 + 10 mM ammonium formate + 0.1% FA. The flow rate was set at 0.3 mL/min and the gradient was as detailed: 0 min 50%B, 1 min 70%B, 6.5 min 100%B, 8.5 min 100%B, 8.6 min 50%B, 9.6 min 50% end run. All gradients were linear.

Mass spectrometry analysis was carried out on a Xevo TQ-XS Triple Quadrupole Mass Spectrometer (Waters Corporation). This was operated in negative ion mode with a source temperature of 150°C and a desolvation gas temperature of 600°C. Nitrogen was used as the desolvation and cone gas at a flow rate of 1000 and 150 L/h, respectively. The capillary voltage was maintained at 3.5 kV, and the cone voltage was set to 30 V. Mass spectral data was analysed using Masslynx software (version 4.2, Waters Corporation). Mass spectra of all CL species were obtained by scanning the m/z range 1000-1600, between 3.5-7 min (scan time 0.2 s). Extracted ion chromatograms (XIC) were then generated for the individual CL species, and the peak areas were integrated. Analyte values were obtained by calculating the ratio of the CL peak areas to the internal standard (C14:0)_4_ CL peak areas. (18:2)_4_ CL, and (C14:0)_4_ CL standards (Avanti Lipids) were used to confirm the retention times of these CL species. Multiple reaction monitoring (MRM), using specific parent and daughter ion transitions, was also used to confirm the identity of these species.

### Statistics

Statistical analysis was performed with GraphPad Prism (version 10.2.2). Data were tested for normality and statistical tests chosen based on normality. If non-parametric, the Mann-Whitney test was used for comparison of two groups. If normally distributed, the unpaired *t*-test was used. Significance was determined as a p value <0.05. Categorical data was analysed by Fishers exact test. Correlation was assessed using the Pearson correlation co-efficient or Spearman rank correlation coefficient depending on normality of data.

## Acknowledgements

This work was supported by intramural funding from the Medical Research Council (MRC) UK to HMC (MC-A654-5QB90) and LF (MC-A654-5QC70). PKM was supported by an NIHR Academic Clinical Fellowship (ACF-2018-21-003), and the Chain Florey Clinician Scientist Training Programme, funded by the MRC (WCMA P64678) and the NIHR Imperial Biomedical Research Centre (WCMA PA6808). MRT is funded by MRC Precision Medicine and the NIHR Imperial BRC. SP was supported by an MRC Clinical Academic Research Partnership grant (MR/V03801X/1).

---

**Supplementary Figure 1.**
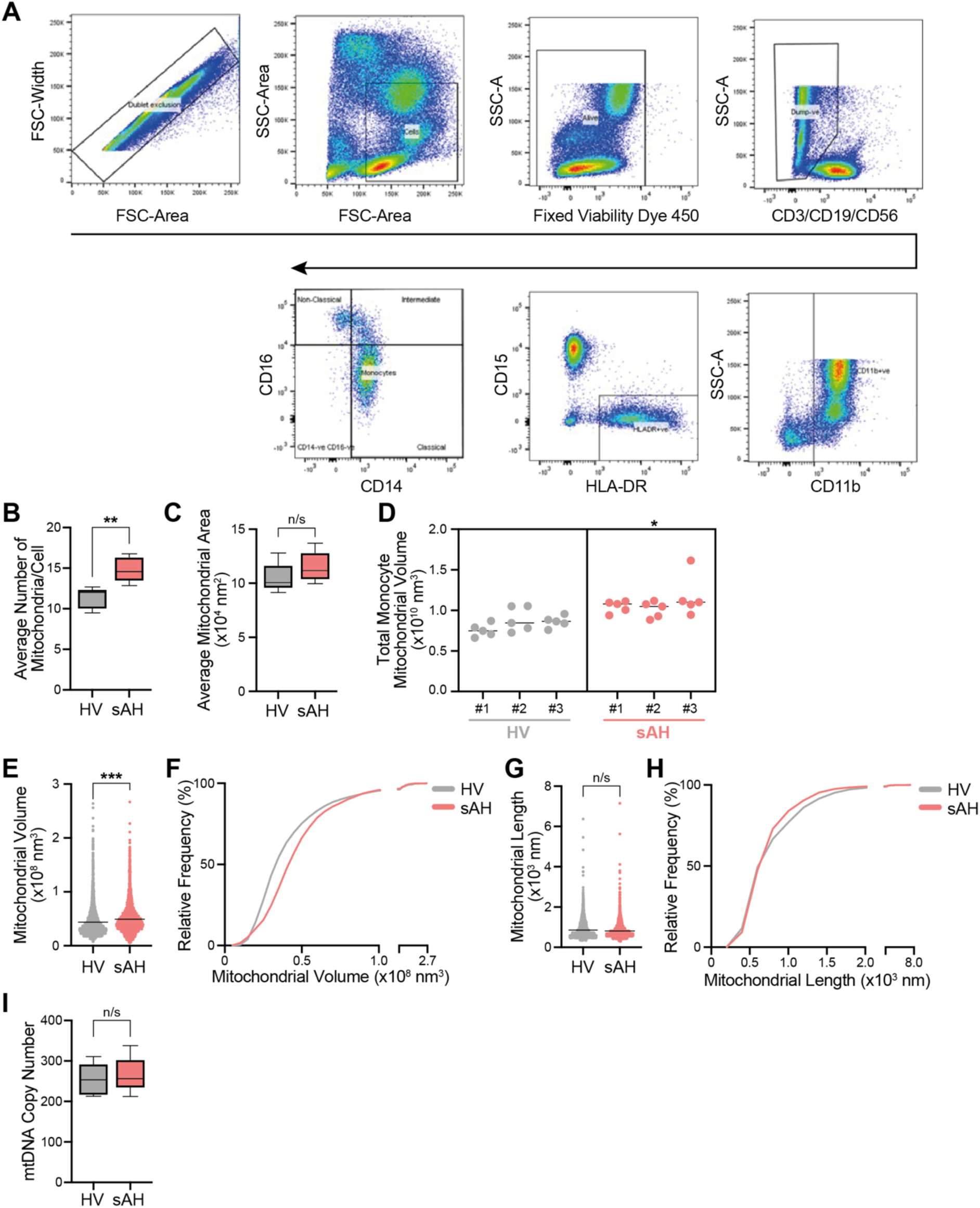
Gating strategy to identify monocytes from leukocytes and additional mitochondrial structural characteristics. **A)** Gating strategy to identify monocytes from lysed whole blood. Monocytes were initially gated as Fixed Viability Dye 450-/CD3-/CD19-/CD56-/CD11b+/HLA-DR+/CD15-cells. Monocyte sub-types were gated by CD14 and CD16 positivity including classical (CD14hi CD16-), intermediate (CD14hi CD16+) and non-classical (CD14lo CD16+). Total monocytes gating included all 3 sub-groups. **B-C)** Analysis of TEM monocyte cross-sections from HV and sAH patients (n=5 individuals, with 25 cells each). Average number of mitochondria (**B**) and average mitochondrial area (**C**). ** p<0.01 (unpaired two-tailed t-test). **D)** Total mitochondrial volume within 5 monocytes from 3 donors. * p<0.05 (nested t-test). **E-H)** SBF-SEM analysis of pooled mitochondria from HV (n=2,844) and sAH (n=3,282). Mitochondrial volume (**E**) and length (**G**), with respective cumulative frequency distributions (**F** and **H)**. n/s, p>0.05, *** p<0.001 (Mann-Whitney test). **I)** Monocyte mtDNA copy number assessed by QPCR (n=5 per condition).

**Supplementary Figure 2.**
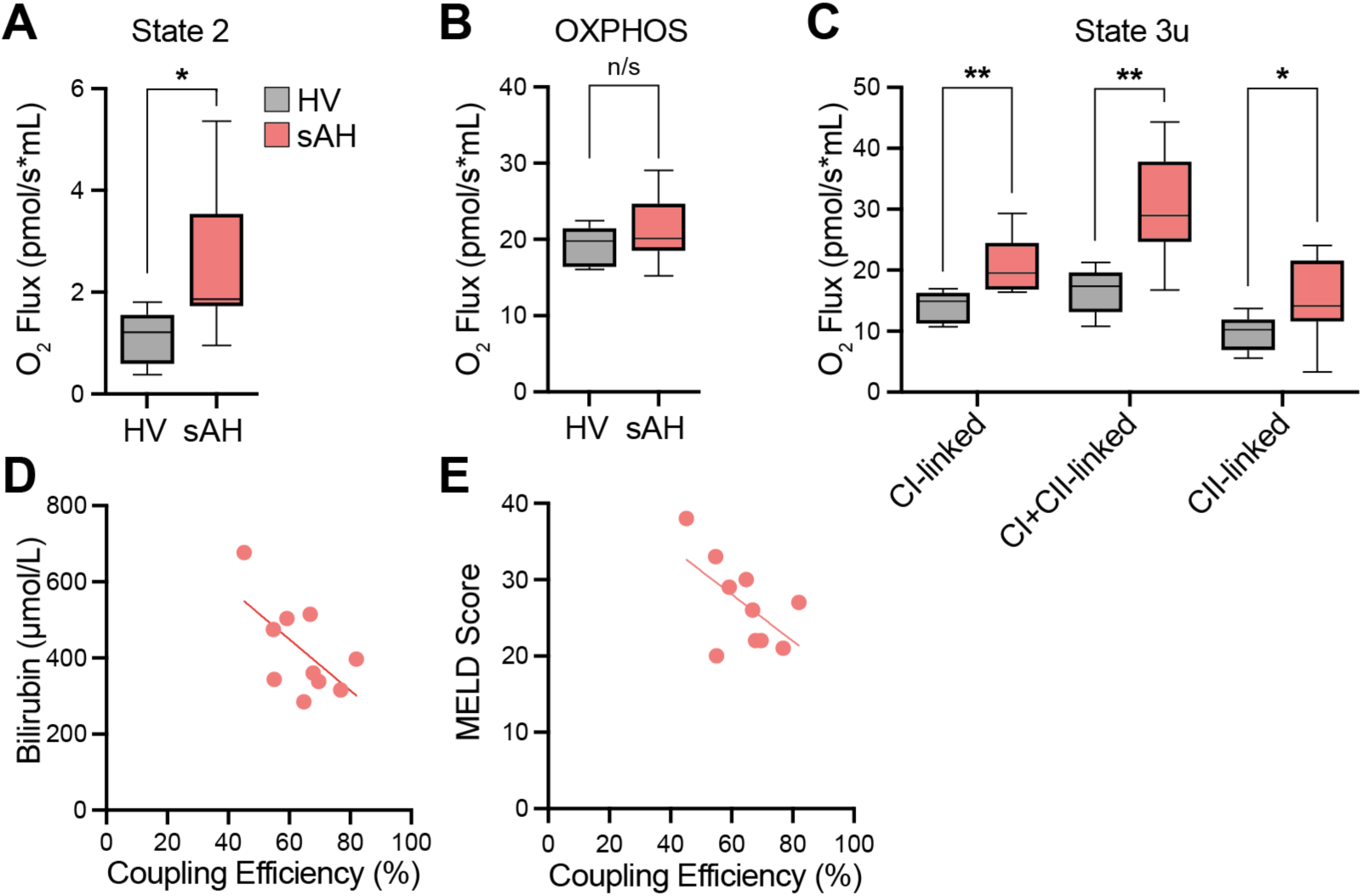
Respirometry on permeabilised monocytes and clinical correlation to coupling efficiency A-C) High resolution respirometry of permeabilised monocytes (1x10^6^ cells) from patients with sAH (n=9) compared to HV (n=5). State 2 respiration (**A**), oxidative phosphorylation capacity (OXPHOS) (**B**), and uncoupled state 3 respiration (State 3u) in the presence of saturating Complex I (CI) and Complex II (CII) substrates (**C**). n/s p>0.05, *p<0.05, ** p<0.01 (Mann-Whitney test). **D-E)** Monocyte mitochondrial respiratory coupling efficiency versus associated bilirubin (**D**) and MELD score (**E**) of sAH patients (n=10) at time of blood draw.

**Supplementary Table 1.**
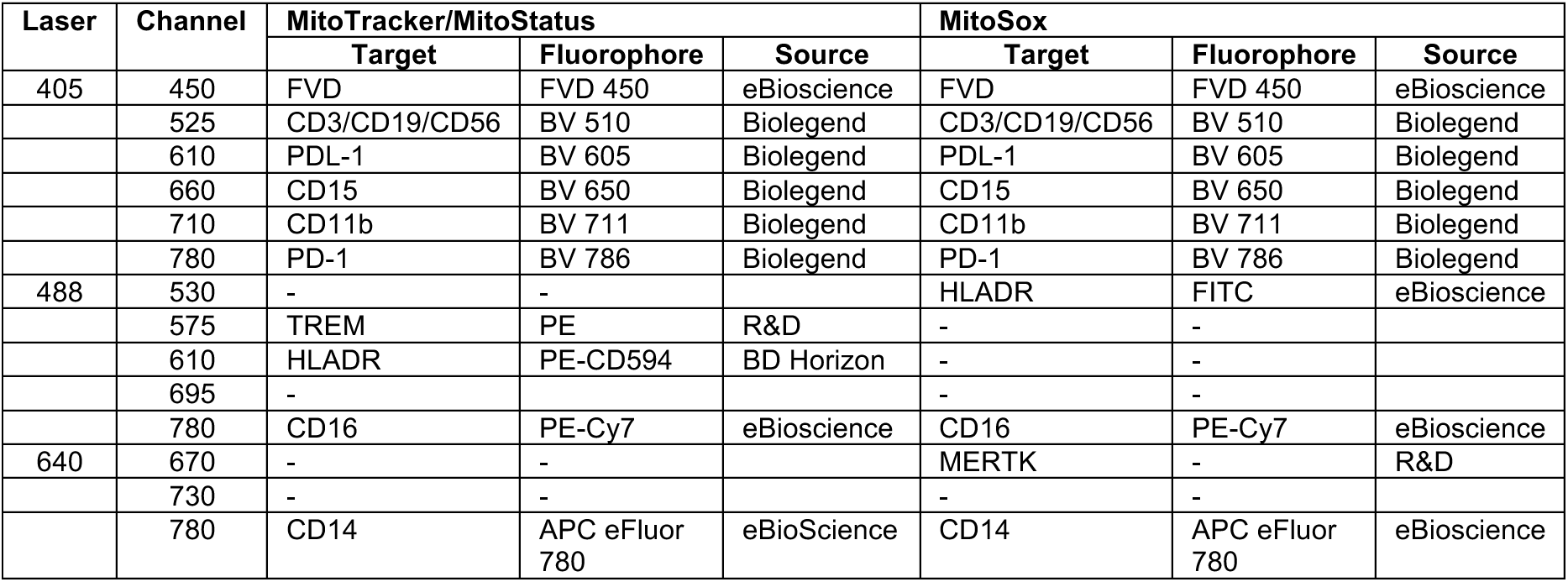
Surface staining antibodies and fluorophores for flow cytometry protocol alongside laser and channel configuration. FVD, Fixed Viability Dye; BV, Brilliant Violet; PDL-1, Programmed Death Ligand 1; CD, Cluster of Differentiation; PD-1, Programmed Cell Death Protein 1; HLA-DR, Human Leukocyte Antigen-DR Isotype; FITC, Fluorescein Isothiocyanate; PE, Phycoerythrin; APC, Allophycocyanin.

**Supplementary Table 2.**
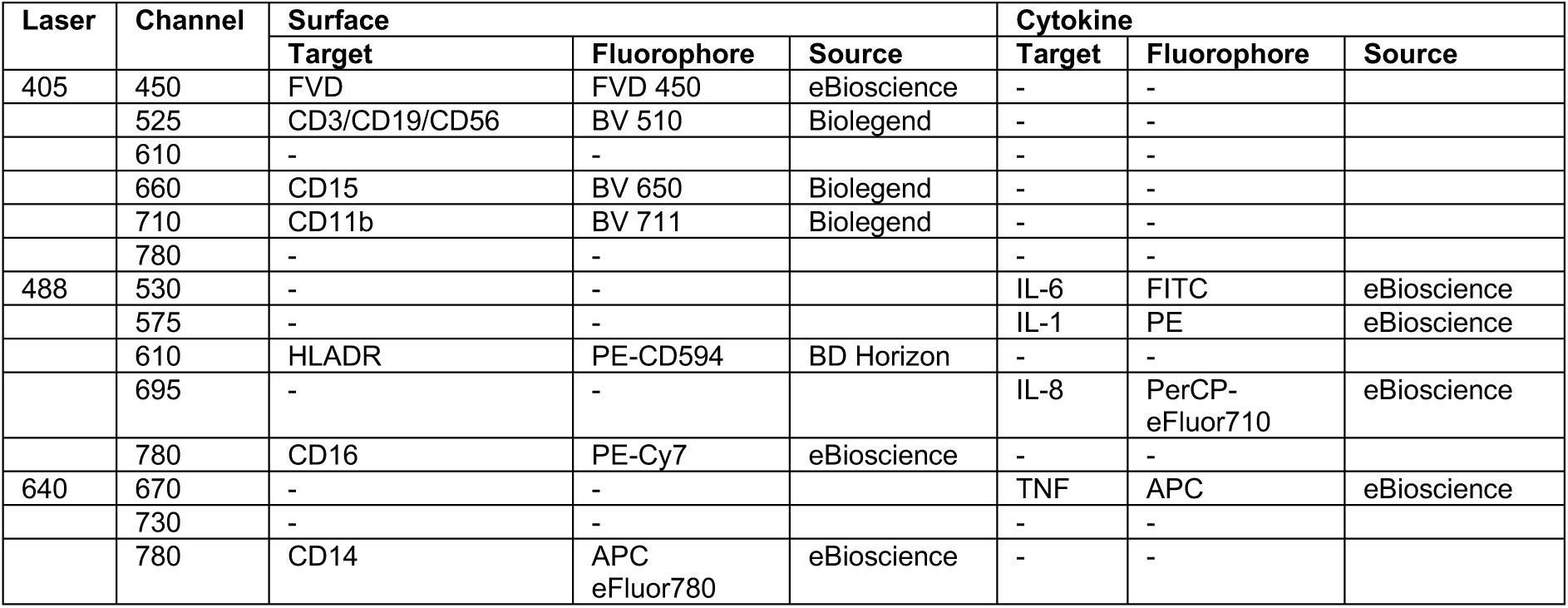
Surface and cytokine antibodies and fluorophores for intracellular cytokine staining flow cytometry protocol alongside laser and channel configuration. FVD, Fixed Viability Dye; BV, Brilliant Violet; PDL-1, Programmed Death Ligand 1; CD, Cluster of Differentiation; PD-1, Programmed Cell Death Protein 1; HLA-DR, Human Leukocyte Antigen-DR Isotype; FITC, Fluorescein Isothiocyanate; PE, Phycoerythrin; APC, Allophycocyanin; IL, interleukin; PerCP, Peridinin-Chlorophyll-Protein.

